# Cytogenetic resource enables mechanistic resolution of changing trends in human pluripotent stem cell aberrations linked to feeder-free culture

**DOI:** 10.1101/2023.09.21.558777

**Authors:** Dylan Stavish, Christopher J. Price, Gabriele Gelezauskaite, Kimberly A. Leonhard, Seth M. Taapken, Erik M. McIntire, Owen Laing, Bethany M. James, Jack J. Riley, Johanna Zerbib, Duncan Baker, Amy L. Harding, Lydia H. Jestice, Thomas F. Eleveld, Ad J.M. Gillis, Sanne Hillenius, Leendert H.J. Looijenga, Paul J. Gokhale, Uri Ben-David, Tenneille E. Ludwig, Ivana Barbaric

## Abstract

Since the first derivation of human pluripotent stem cells (hPSCs), the number of culture conditions has steadily increased, making hPSC culture more facile. Nonetheless, there remains the persistent issue of culture-acquired genetic changes, hampering the reproducibility of hPSC research and jeopardising their clinical use. Here, we utilised comprehensive karyotyping datasets from over 20,000 hPSC cultures sampled under different conditions to ascertain association of genetic changes with specific culture regimens. We found condition-dependent patterns of aberrations, with higher prevalence of chromosome 1q gains in recent years, associated with increased use of contemporary, feeder-free cultures. Mechanistically, we show the context-dependent selection of 1q variants is mainly driven by *MDM4*, a gene amplified in many cancers, located on chromosome 1q. To facilitate reproducibility of hPSC research and their safe clinical utility, we provide a unique hPSC karyotype resource for informing the risk assessment of genetic aberrations and developing strategies to suppress their occurrence.

## Introduction

The dual ability of human pluripotent stem cells (hPSCs) to self-renew seemingly indefinitely *in vitro* whilst remaining capable of trilineage differentiation has made them a potent tool in basic research and regenerative medicine^1^. However, the occurrence of culture-acquired genetic changes in hPSCs can confound experimental results and stands to jeopardise the progress of hPSC-based cell replacement therapies, as genetic changes may impact on growth properties, differentiation ability and tumorigenic potential of hPSCs^2–4^. In line with these observations, the recently-developed International Society for Stem Cell Research (ISSCR) guidelines for the use of stem cells in research (ISSCR Standards; https://www.isscr.org/standards) stipulate assessing genetic integrity of hPSC lines as a measure towards improving the rigour and reproducibility of data in the field^5^. Yet, as highlighted in ISSCR Standards and various reviews^2,4,6^, the challenges remain in understanding which specific changes to monitor and how to interpret their functional consequences.

Although genetic changes in hPSCs are known to occur at all scales, from single nucleotide variants to karyotypic abnormalities, on the whole, karyotypic changes may be expected to have greater detectable phenotypic effect due to a large number of genes typically implicated. Karyotyping of different hPSC lines from independent cultures and different laboratories across the world, has led to the conclusion that the culture-acquired chromosomal aberrations in hPSCs exhibit a recurrent pattern, with gains of whole or parts of chromosomes 1, 12, 17, 20 and X being particularly prevalent^7–13^. Since hPSCs were first derived on fibroblast feeder layers and in the presence of Knock-Out Serum Replacement (KOSR)^14^, a range of feeder-free/KOSR-free conditions became available. Hence, a question arises which, if any, of the existing conditions afford better karyotypic stability of hPSCs? Moreover, for purposes of monitoring the expanding hPSC cultures, critical missing information is whether recurrent genetic abnormalities differ depending on the culture conditions. Finally, strategies for suppressing variants require identification of driver gene(s) and mechanisms underpinning the growth advantage of variant cells.

To address these questions, we used karyotyping data from over 20,000 samples to provide an up-to-date overview of chromosomal regions subject to frequent alterations in hPSCs and to reveal how changes in culture conditions impact patterns of recurrent aberrations. We further identify a gene on chromosome 1q, the TP53 inhibitor *MDM4,* as giving the selective advantage to hPSCs grown in contemporary, feeder-free/KOSR-free culture conditions.

## Results

### Changes in aneuploidy patterns in hPSCs over time

We reasoned that a retrospective analysis of a large dataset of curated karyotypes with associated culture conditions metadata would enable us to make correlations and generate specific hypotheses regarding karyotypic abnormalities associated with particular culture regimens that could be empirically tested and mechanistically investigated. To this end, we analysed karyotypic aberrations from a large dataset of 19,942 hPSC karyotypes acquired by WiCell (www.wicell.org) between 2009 and 2021, the majority of which have been annotated with features including the date of the cytogenetic analysis and culture conditions used (**Figure 1A; Table S1)**. In tandem, we also analysed a smaller in-house dataset from the Centre for Stem Cell Biology (CSCB) in Sheffield, containing 1,772 karyotypes collected between 2002-2019 (**Figure 1A; Table S1**). We confirmed the compatibility of the datasets by examining the occurrence and types of abnormalities recorded. In both datasets abnormal clones were present at a similar percentage, i.e., 25% and 22% for WiCell and CSCB, respectively (**Figure 1B**). Additionally, in both datasets gains of chromosomes or chromosomal regions were the most common karyotypic aberration (**Figure 1C**). This was followed by gains and losses of chromosomal material (**Figure 1C**), the majority of which manifested as isochromosomes (**Figure 1D**). Aberrations that entailed only the loss of chromosomes or parts of chromosomes were relatively infrequent, as were balanced translocations (**Figure 1C)**. Overall, in line with previous reports^9–11,13^, this data showed that the appearance of karyotypically abnormal cells is a relatively frequent occurrence in hPSC cultures, with gains of chromosomal materials being more prevalent than losses.

**Figure 1.**
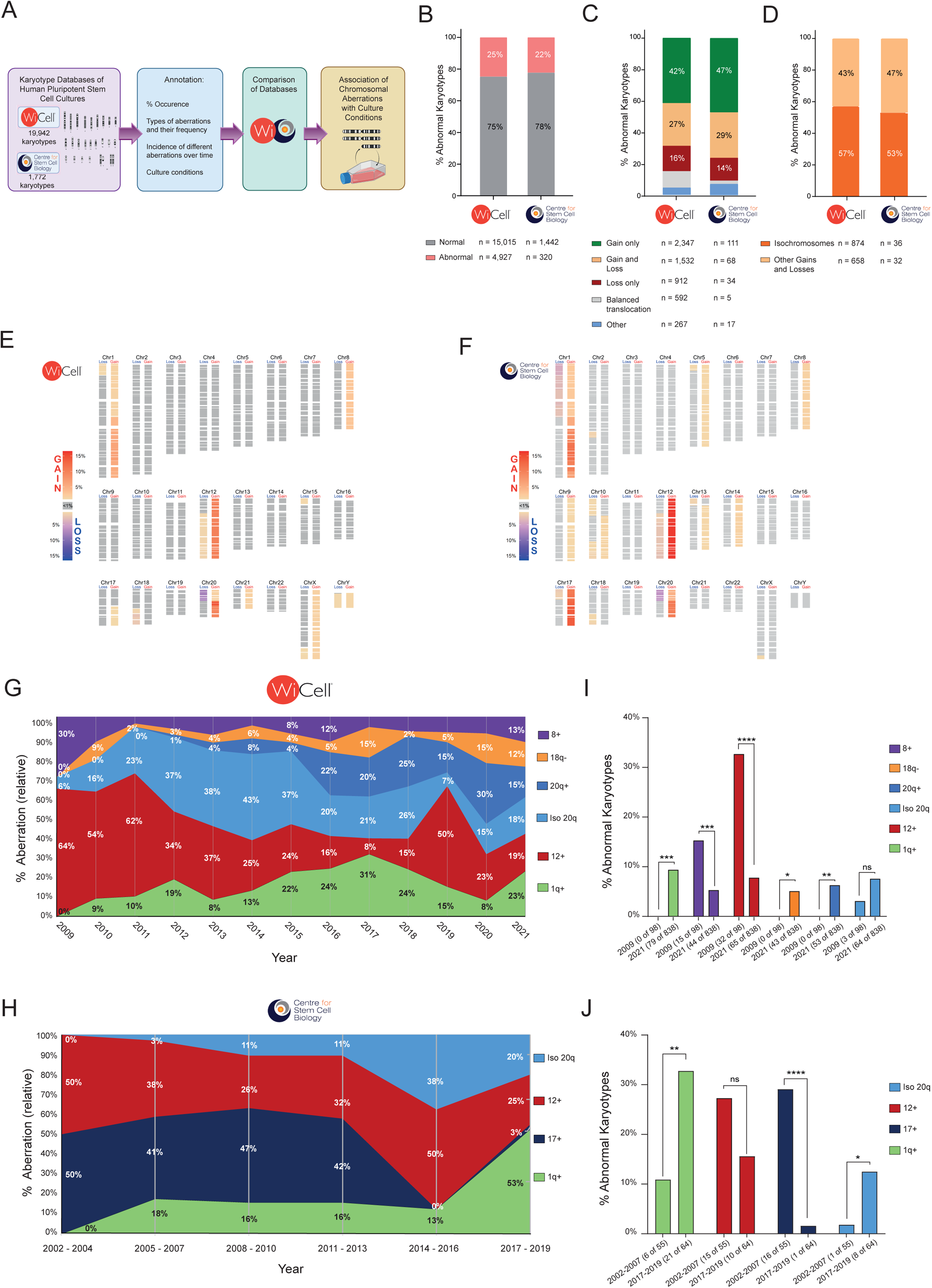
Retrospective analysis of aberrant karyotypes and conditions for culturing hPSCs. **(A)** Karyotyping data from two independent centres (WiCell and Centre for Stem Cell Biology (CSCB)) were analysed and annotated for different parameters, including date of collection and culture conditions. **(B)** Percentage of hPSC cultures containing cells with abnormal karyotypes is similar between WiCell and CSCB datasets. **(C)** The breakdown of abnormal karyotypes according to the type of abnormality. **(D)** The majority of aberrations involving both gains and losses of chromosomal material are isochromosomes. **(E)** The frequency of gains/losses of each cytoband across all abnormal karyotypes in the WiCell dataset. **(F)** The frequency of gains/losses of each cytoband across all abnormal karyotypes in the CSCB dataset. **(G)** The relative frequency of the six most common abnormalities in the WiCell dataset over time. **(H)** The relative frequency of the four most common abnormalities in the CSCB dataset over time. **(I)** The proportion of abnormal karyotypes containing one of the six most common abnormalities in the WiCell dataset sampled in the year 2009 versus the year 2021. ns, non-significant; *p<0.05, **p<0.01, ***p<0.001, ****p<0.0001; Fisher’s Exact test. **(J)** The proportion of abnormal karyotypes containing one of the four most common abnormalities in the CSCB dataset sampled in 2002-2007 versus 2017-2019 (several years are pooled together due to relatively low n numbers in this dataset). ns, non-significant; *p<0.05, **p<0.01, ****p<0.0001; Fisher’s Exact test. See also **Table S1**, **Figure S1**, **S2**.

To identify recurrent abnormalities in each dataset we used CytoGPS^15^, which converts traditional karyotype nomenclature into gain and loss projections by G-band (**Table S2**). This analysis confirmed a non-random distribution of chromosomal aberrations in each dataset (**Figure 1E,F; Figure S1, S2**). Next, we defined the recurrent abnormalities as clonal aberrations that were present in more than 1% of the total karyotypes. While the similarities between the datasets included high frequency of abnormal karyotypes with gains of chromosome 1q (9.3% in WiCell; 14% in CSCB), trisomy 12 (13.7% in WiCell; 20% in CSCB) and gains of isochromosome 20q (7.8% in WiCell; 8.1% in CSCB) (**Table S2**), we also noted some differences. For instance, 18q loss and trisomy 8 that were frequent in the WiCell dataset (3.9% and 4.6%, respectively) were present in less than 1% of total karyotypes in the CSCB dataset (**Table S2**). In contrast, the CSCB dataset contained a high frequency of trisomy 17 (14.5%), whilst in the WiCell data this aberration represented only 0.6% of all abnormalities (**Table S2**).

Next, we assessed whether the incidence of the most frequently encountered aberrations in WiCell and CSCB datasets was constant with time. Strikingly, we found a disparity in the representation of different variants over time (**Figure 1G,H**), with an overall trend towards increase in frequency of aberrations such as gains of chromosome 1q, 20q and isochromosome 20q at the expense of aberrations such as trisomy 12 and 17 (**Figure 1G-J**). In summary, patterns of karyotypic abnormalities have changed over the last two decades, with 20q and 1q gains becoming more prevalent – and trisomy 12 and 17 becoming less prevalent - in recent years.

### Increased frequency in chromosome 1q gains correlates with usage of KOSR-free culture conditions

Over time, culture conditions have shifted from fibroblast feeders and KOSR-containing media (from herein termed KOSR-based) to a range of feeder-free, KOSR-free regimens (from herein termed KOSR-free) (**Figure 2A,B)**. We therefore reasoned that the observed changes in variant frequency may be due to these changes in culture conditions. To test this hypothesis, we used our annotated karyotyping datasets to assess a potential association of aberrant karyotypes with specific culture conditions. We first assessed whether the percentage of abnormal karyotypes differed significantly depending on the culture medium or matrix used. We found a similar percentage (18-22%) of abnormal karyotypes in the KOSR-based and four most represented KOSR-free media across the WiCell dataset (E8, mTeSR, NutriStem and StemFlex) (**Figure 2C**). This was also true for the six most used matrices in the WiCell data (feeders, geltrex, matrigel, laminin 511, laminin 521 and vitronectin) (**Figure 2D**). Together, this analysis indicated that the overall incidence of karyotypic changes is similar across culture media and matrices.

**Figure 2.**
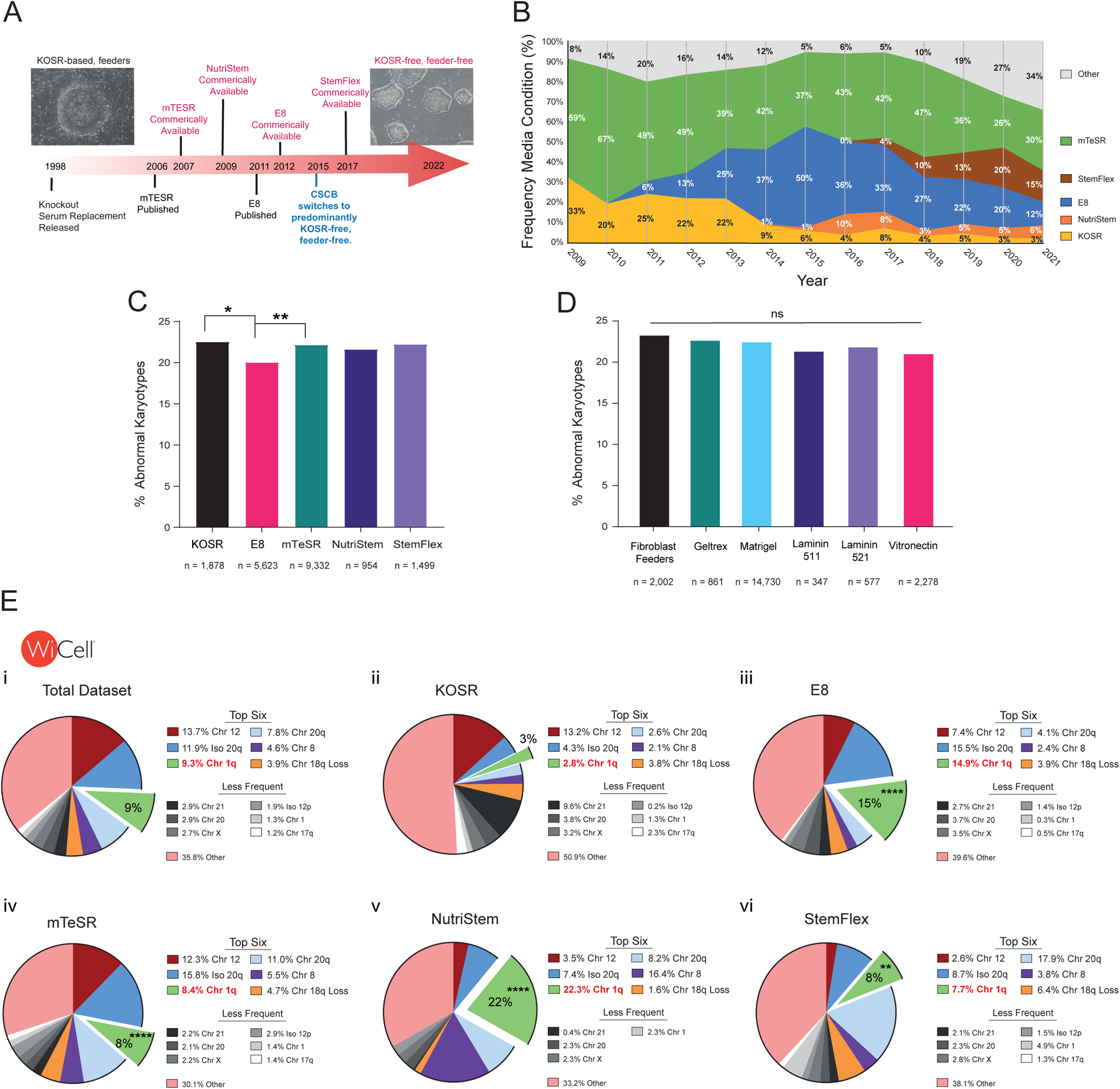
Gains in chromosome 1q are associated with KOSR-free conditions. **(A)** Timeline of the introduction of popular hPSC culture conditions over the past 25 years beginning with predominantly KOSR-based systems and transitioning into KOSR-free culture regimens. CSCB started using predominantly KOSR-free systems in 2015. **(B)** The relative frequency of the most used media in the WiCell dataset over time. The use of KOSR-based media had decreased over time. **(C)** The percentage of abnormal karyotypes from the cultures using the most represented media across the WiCell dataset. *p<0.05; **p<0.01; Fisher’s Exact test. **(D)** The percentage of abnormal karyotypes from the cultures using the most represented matrices across the WiCell dataset. ns, non-significant; Fisher’s Exact test. **(E)** The percentage of the most common abnormalities in the WiCell dataset across (i) all media, (ii) KOSR-based media, (iii) E8, (iv) mTESR, (v) NutriStem and (vi) StemFlex. Less frequent but recurrent aberrations (above 1% of abnormal karyotypes) are indicated in shades of grey. In comparison to KOSR-based medium, the frequency of 1q gain is elevated in E8 (****p=7.6×10^−15^), mTESR (****p=4.5×10^−6^), NutriStem (****p=8.9×10^−17^) and StemFlex (**p=0.0014); Fisher’s Exact test.

We next asked whether the identity of the chromosome involved in abnormal karyotypes differed based on the medium in which the cells were grown. We assessed the recurrent abnormality landscape for each of the most represented media in the WiCell dataset and found that the relative frequency of commonly acquired chromosomal aberrations differed across different conditions (**Figure 2Ei-vi**). Of note, while gains of chromosome 1q represented the third most common abnormality (∼9% of all abnormal karyotypes) across the entire dataset (**Figure 2Ei**), this aberration was over-represented in the KOSR-free media in comparison to the KOSR-based conditions (8-22% versus 3%, respectively) (**Figure 2Eii-vi**). Together, our analysis revealed that the landscape of common abnormalities in hPSCs had changed over time, seemingly coinciding with changes in culture conditions utilised. Specifically, we detected an increase in the prevalence of chromosome 1q gains in association with the usage of KOSR-free conditions in recent years.

### Variant hPSCs with a gain of chromosome 1q show selective advantage in a context-dependent manner

We posited that the increased frequency of chromosome 1q gains amongst karyotypes of hPSCs grown in KOSR-free conditions is the result of context-dependent selection, with 1q variant cells possessing the selective advantage in KOSR-free, but not KOSR-based conditions. To test this hypothesis, we utilised pairs of cells with a gain of a portion of chromosome 1q (from herein *v1q*) and their isogenic wild-type counterparts across several different genetic backgrounds: H7 (WA07), H9 (WA09), WLS-1C and MIFF-3 (**Figure 3A**; **Figure S3**). Of note, the 1q gain in the MIFF-3 *v1q* line was only detected by SNP-arrays, as the amplified region was below the resolution of G-banding (**Figure S3**).

**Figure 3.**
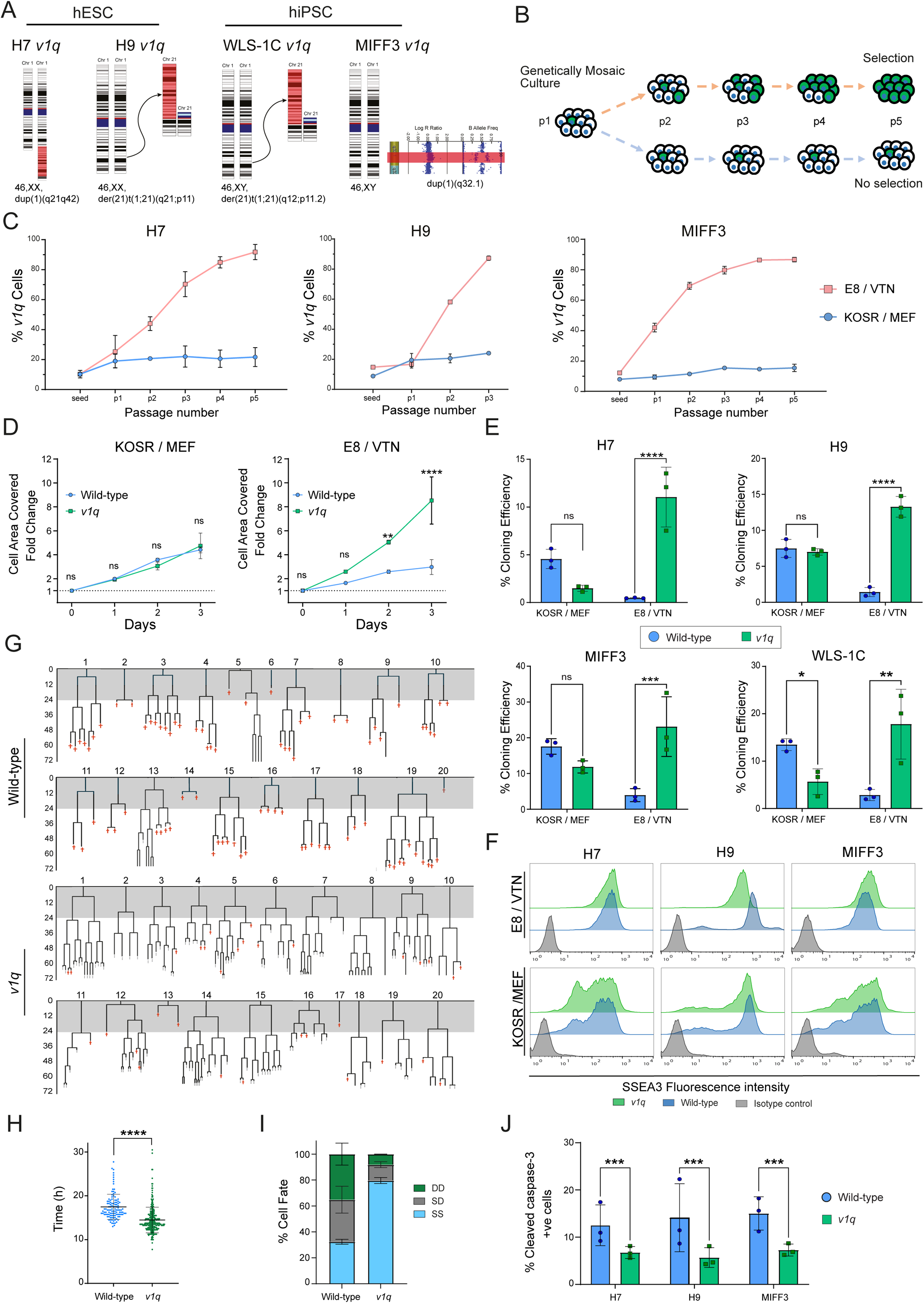
*v1q* have selective advantage in KOSR-free but not KOSR-based conditions. **A.** A panel of wild-type and *v1q* sublines across four genetic backgrounds (H7, H9, WLS-1C and MIFF3) used in this study. **B.** Selective advantage was tested by mixing ∼10% *v1q* with their wild-type counterparts, with either of the lines being fluorescently labelled. Mixed cells were plated into either KOSR/MEF or E8/VTN and the ratio of variants was monitored over subsequent passages. **C.** *v1q* overtake wild-type cells rapidly in E8/VTN but not in KOSR/MEF. Data shown are the mean ± SD of three biological replicates. **D.** *v1q* have a significantly higher growth rate than wild-type cells in E8/VTN but not KOSR/MEF. Data shown are the mean ± SD of three biological replicates. ns, non-significant; **p<0.01, ****p<0.0001; Two-way ANOVA. **E.** *v1q* have a significantly higher cloning efficiency than wild-type cells in E8/VTN but not KOSR/MEF. Data shown are the mean ± SD of three biological replicates. ns, non-significant; *p<0.05, **p<0.01, ***p<0.001, ****p<0.0001. Two-way ANOVA, Fisher’s LSD. **F.** Wild-type and *v1q* hPSCs display similar levels of expression of a marker of undifferentiated state, SSEA3, in E8/VTN and in KOSR/MEF conditions. **G.** Lineage trees tracked from time-lapse images of wild-type cells (upper two panels) and *v1q* (lower two panels). Red crosses indicate cell death. Gray-shaded area indicates the first 24 h post-plating when the cells were grown in the presence of Y-27632, required for single cell passaging. **H.** *v1q* display faster cell cycle compared to wild-type counterparts. Data points indicate 114 and 286 divisions for wild-type and *v1q* cells, respectively, from two independent experiments. ****p<0.0001; Mann-Whitney test. **I.** Percentage of cell fate outcomes of daughter cells following cell division, with SS denoting survival of both daughter cells, SD survival of one and death of the other daughter cell and DD death of both daughter cells. **J.** *v1q* have decreased levels of cleaved caspase-3 marker of apoptosis. Data shown are the mean ± SD of three biological replicates. ***p<0.001; Two-way ANOVA followed by Holm-Sidak’s multiple comparison test. See also **Figure S3, S4**; **Video S1**.

We first performed competition experiments in which we mixed a small proportion (∼10%) of *v1q* cells into wild-type cultures and then monitored the ratio of the two sublines over subsequent passages either in KOSR-based medium on mouse embryonic fibroblasts (KOSR/MEF) or in KOSR-free E8/vitronectin (E8/VTN) conditions (**Figure 3B**). Strikingly, whilst in KOSR-based conditions, the ratio of *v1q* overall remained unchanged over several consecutive passages, in E8/VTN *v1q* rapidly overtook the cultures (**Figure 3C**). *v1q* cells also displayed faster population growth in comparison to wild-type cells only in E8/VTN, but not in KOSR/MEF condition (**Figure 3D**). Finally, we utilised a clonogenic assay as a particularly sensitive test of the ability of individual hPSCs to survive replating and initiate stem cell colonies^16^. The clonogenic assay showed an increased cloning ability of *v1q* compared to wild-type hPSCs, again only when cloning was performed in E8/VTN, but not KOSR/MEF conditions (**Figure 3E**). Overall, these data support the hypothesis that *v1q* hPSCs have a selective advantage over wild-type hPSCs in E8/VTN, but not KOSR-based cultures. Importantly, we found no evidence of differences in the levels of spontaneous differentiation between paired wild-type cells and *v1q* (**Figure 3F; Figure S4A**), suggesting that differences in survival and/or proliferation, rather than differences in the level of spontaneous differentiation, underpin the selective advantage of *v1q*. Indeed, time-lapse analysis of single cells revealed a shorter cell cycle time of *v1q* (**Figure 3G,H; Video S1; Figure S4B)** and their improved survival, as fewer numbers of *v1q* underwent cell death upon replating and following cell division compared to their wild-type counterparts (**Figure 3G,I; Video S1; Figure S4B**). Consistent with these findings, a lower proportion of cells in *v1q* cultures displayed the cleaved caspase-3 marker of apoptosis (**Figure 3J**). Together, these results reveal that increased proliferation and reduced apoptosis of *v1q* cells underscore their selective advantage in E8/VTN condition.

### MDM4 drives the selective advantage of 1q variant hPSCs in E8/VTN condition

To identify potential driver genes underpinning the selective advantage of a gain of chromosome 1q in E8/VTN, we mapped the minimal region amplified amongst all the *v1q* in the karyotyping datasets (**Figure 4A**). Further, overlaying the SNParray data of the MIFF-3 *v1q* line with a previously-reported 1q minimal amplicon in hPSCs^17^ enabled us to narrow the candidate region on chromosome 1q32.1 to ∼1Mb, encompassing 13 genes expressed in hPSCs (**Figure 4B**). We ranked the expressed genes based on their essentiality to the pluripotent state^18^ (**Figure 4B**). Notably, one of the top three most essential genes within this region is *MDM4*, a known regulator of TP53^19^ and a putative driver gene in the pathology of multiple types of cancers^20,21^. RNA-seq analysis of H7 and H9 *v1q* versus wild-type sublines further revealed differential expression of the p53 pathway in the *v1q* hPSCs (**Figure 4C**). Thus, we posited that the increase in copy number of *MDM4* due to the amplification of chromosome 1q32 could provide selective advantage of *v1q* in E8/VTN conditions.

**Figure 4.**
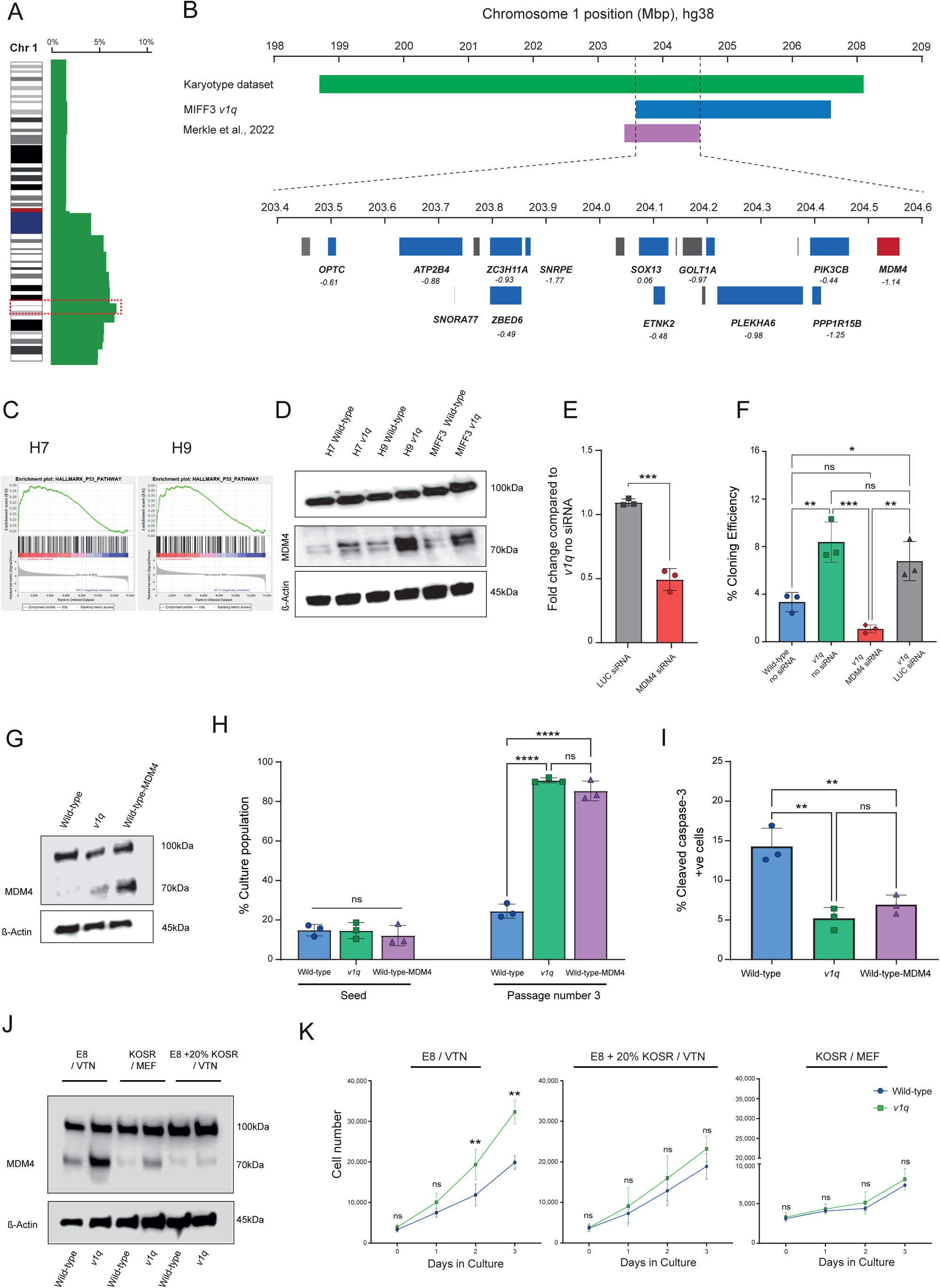
MDM4 overexpression provides selective advantage to *v1q*. **A.** The minimal region on chromosome 1q32.1 identified from the karyotyping datasets in this study. **B.** The minimal amplicon contains 13 genes expressed in hPSCs (blue boxes), which were ranked based on their essentiality scores (indicated in italics underneath the genes). *MDM4* (red) is a candidate of interest, based on its known role in p53 signalling and cancer. **C.** RNA-seq analysis of H7 and H9 *v1q* versus wild-type sublines revealed differential expression of the p53 pathway. **D.** MDM4 expression is increased in *v1q*. Western blot analysis of H7, H9 and MIFF3 wild-type and v1q cells. β-actin was used as a loading control. **E.** Knockdown of MDM4 with siRNA in *v1q* was confirmed by quantitative PCR. siRNA for Renilla Luciferase (siRNA LUC) was used as a negative control. Data shown are the mean ± SD of three biological replicates. ***p<0.001; Student’s t test. **F.** MDM4 knock-down suppresses cloning efficiency of *v1q*. Data shown are the mean ± SD of three biological replicates. ns, non-significant, *p<0.05, **p<0.01, ***p<0.001; One-way ANOVA followed by Holm-Sidak’s multiple comparison test. **G.** Overexpression of MDM4 in wild-type hPSCs. Western blot analysis of MIFF3, MIFF3 *v1q* and MIFF3 wild-type cells overexpressing MDM4 (Wild-type-MDM4). β-actin was used as a loading control. **H.** MDM4 overexpression provides selective advantage to wild-type hPSCs. Mixed cultures were analysed at seeding and after 3 passages. Data shown are the mean ± SD of three biological replicates. ns, non-significant, ***p<0.001; One-way ANOVA followed by Holm-Sidak’s multiple comparison test. **I.** MDM4 overexpressing and *v1q* cells have lower percentage of cleaved caspase-3 marker of apoptosis compared to wild-type counterparts. Data shown are the mean ± SD of three biological replicates. ns, non-significant, **p<0.01; One-way ANOVA followed by Holm-Sidak’s multiple comparison test. **J.** MDM4 abundance is reduced in KOSR/MEF. Also, the addition of KOSR to E8/VTN reduces MDM4 expression. Western blot analysis of MDM4 in MIFF3 and MIFF3 *v1q* cells under different conditions. β-actin was used as a loading control. **K.** The addition of KOSR to E8/VTN reduces growth rates of *v1q* to levels like those of wild-type cells. Data shown are the mean ± SD of three biological replicates. ns, non-significant; *p<0.05, **p<0.01; Two-way ANOVA with Holm-Sidak’s multiple comparison test.

Indeed, in E8/VTN conditions, *v1q* showed higher abundance of MDM4 protein expression, consistent with an increased copy number of *MDM4* (**Figure 4D**). To functionally probe the role of MDM4 in hPSC selective advantage, we knocked-down MDM4 in *v1q* cells (**Figure 4E)**, which resulted in significant decrease in their cloning ability (**Figure 4F**). Conversely, we overexpressed MDM4 in wild-type hPSCs and found that increase in MDM4 (**Figure 4G**) phenocopied the selective advantage of *v1q*, as MDM4 overexpressing cells outcompeted wild-type cells in competition experiments, to the same extent as *v1q* did (**Figure 4H**). Moreover, MDM4 overexpressing cells exhibited lower levels of cleaved caspase-3 compared to wild-type cells (and similar levels to those of *v1q*; **Figure 4I**), consistent with the acquisition of an anti-apoptotic phenotype due to MDM4 overexpression. Together, these data demonstrate that MDM4 amplification is a key contributor to the selective advantage of *v1q* in E8/VTN conditions.

Why is the MDM4-mediated 1q gain effect specific to the E8/VTN condition? In the KOSR/MEF condition, we noted reduced abundance of MDM4 in the *v1q* cells compared to E8/VTN conditions (**Figure 4J**). Further, the addition of KOSR to E8/VTN cultures abolished MDM4 expression (**Figure 4J**) and suppressed proliferation rates of *v1q* (**Figure 4K**). Overall, consistent with the context-dependent advantage of variant cells, these data suggest that KOSR/MEF conditions somehow reduce MDM4 protein expression, thereby drastically attenuating the selective advantage of *v1q* hPSCs. A clear implication of these findings is that the recurrent genetic changes observed in hPSCs are directly related to the selection pressures conferred by their *in vitro* environment.

### KaryoBrowser Tool

Currently, many of the detected hPSC genetic aberrations are not reported in literature, thus preventing the identification of potential secular trends in their appearance. To address this issue and share the knowledge of genetic changes arising in hPSCs, we provide here an explorative browser-based database termed KaryoBrowser (**Figure S5**). Using KaryoBrowser, the collated datasets in this study can be inspected and filtered based on different parameters, including culture media, matrix, whether the lines are embryonic or induced PSCs and their sex. Interactive ideograms can be generated as well as bar charts detailing the chromosomal alterations associated with each parameter. The filtered data containing the karyotypes can also be exported for further analysis.

## Discussion

The recurrent nature of culture-acquired genetic changes in hPSCs has been attributed to their ability to confer growth advantage to variant cells^22^, reminiscent of recurrent patterns of aneuploidy seen in most cancers^23^. Culture conditions for hPSCs have evolved and diversified over the last two decades, raising a possibility that different genetic changes may be selected for under different culture conditions. Tracking and correlating particular abnormalities with culture regimens is difficult in a classic laboratory setting due to low sample numbers and a narrow range of conditions typically assessed. Here, we have demonstrated how a large dataset, sampled across two decades from multiple sources and conditions allows for secular trends in culture-acquired genetic changes to be identified and mechanistically probed.

Consistent with previous reports^9–11^, we found that a large proportion (75-78%) of hPSC karyotypes appeared as normal diploid, attesting to the fact that hPSCs are not inherently genetically unstable^24^. Still, the fact that almost one in four of hPSC cultures contains karyotypically aberrant clones is far from ideal for applications of these cells in research and medicine. Interestingly, the proportion of abnormal karyotypes was relatively similar across a range of commonly used media and matrix conditions, suggesting that, from the genetic stability perspective, none of the tested conditions are ideal for long-term hPSC maintenance. Strikingly, despite yielding an overall similar proportion of abnormal karyotypes, different culture conditions evidently selected for different types of aberrations. A case in point is a gain of chromosome 1q, which appeared more prevalent in KOSR-free compared to KOSR-based conditions. Timeline analysis of karyotypic aberrations also showed a recent rise in the prevalence of chromosome 1q gains, coinciding with the increased usage of KOSR-free culture regimens. Utilising hPSC lines paired for the presence or absence of chromosome 1q gain in competition experiments, we confirmed that *v1q* confers proliferation advantage under KOSR-free (E8/VTN), but not KOSR-based conditions.

To identify a driver gene on chromosome 1q, we focused on a p53 inhibitor *MDM4* located in the minimally amplified region. *TP53* was previously demonstrated to be the most growth-restricting^18^ and also the most frequently mutated gene in hPSCs^25,26^, attesting to the importance of p53 in hPSC biology and suggesting that regulators of p53, such as MDM4, may also be a key target of genetic changes. Here, we showed that increased expression of MDM4 in wild-type hPSCs phenocopies *v1q*, suggesting that *MDM4* is a likely driver gene for this amplification in hPSCs. Nonetheless, it is important to note that additional genes present in the amplified region of chromosome 1q could also play a role in the behaviour of variant cells. Intriguingly, chromosome 1q is amplified in many cancers^19^ and *MDM4* was recently proposed as a driver of recurrent chromosome 1q gains in breast cancer through increased expression of MDM4 and reduced TP53 signalling^21^. These parallels between variant hPSCs and cancer reinforce concerns regarding their safety for regenerative medicine.

Understanding of the relationship between genetic changes and conditions under which they are selected for is important not only for prioritising variants to monitor under specific conditions, but also for devising strategies for minimising their appearance. Nonetheless, most of the abnormalities noted in routine hPSC karyotyping are not reported in literature, making up-to-date findings on abnormalities inaccessible to the community. We propose the need for a database collating genetic changes in hPSCs to enable timely identification of links between culture practices and genetic aberrations. As a starting point, we provide a browser-based database of aberrations and their associated metadata analysed in this study.

Overall, our discoveries set the stage for approaches to the detection of genetic changes in hPSCs and targeted strategies for minimising the appearance of recurrent aberrations.

### Limitations of the study

The first part of our study was a retrospective analysis that may have been affected by ascertainment bias, e.g., we do not assert that these are different cell lines, unrelated samples or considered normal at the time of testing. Also, the culture medium used in the dataset was the medium in which the line was submitted in for karyotyping and not necessarily the only culture platform it had been grown in. While we empirically validated association of *v1q* with E8/VTN culture system, further studies are required to confirm other associations of genetic changes with particular culture regimens or different parameters uncovered from the dataset. Given the need for monitoring genetic changes in hPSCs used for basic research or clinical applications, prospective analyses with a more careful curation of genetic abnormalities will allow a timely identification of links between hPSC culture practices and genetic aberrations.

## SUPPLEMENTAL INFORMATION

Supplemental Information includes 5 figures, 2 tables and 1 video.

## Acknowledgments

This work was supported by MR/R015724/1, MR/X000028/1, MR/X007979/1, BB/M011151/1, BB/T007222/1 and MRC DiMeN studentships. We are grateful to Nigel N. Khan for help with developing KaryoBrowser. The WLS-1C *v1q* cells were kindly provided by STEMCELL Technologies. The parental WLC-1C line was generated by Ottawa Human Pluripotent Stem Cell Facility.

## Conflicts of interest

T.L. is a co-inventor and receives a share of royalties on various hPSC media- and culture-related patents currently owned and licensed by the Wisconsin Alumni Research Foundation (WARF). I.B. is a member of the scientific advisory board of WiCell.

## SUPPLEMENTAL INFORMATION

### Supplemental Figure Legends

**Figure S1.**
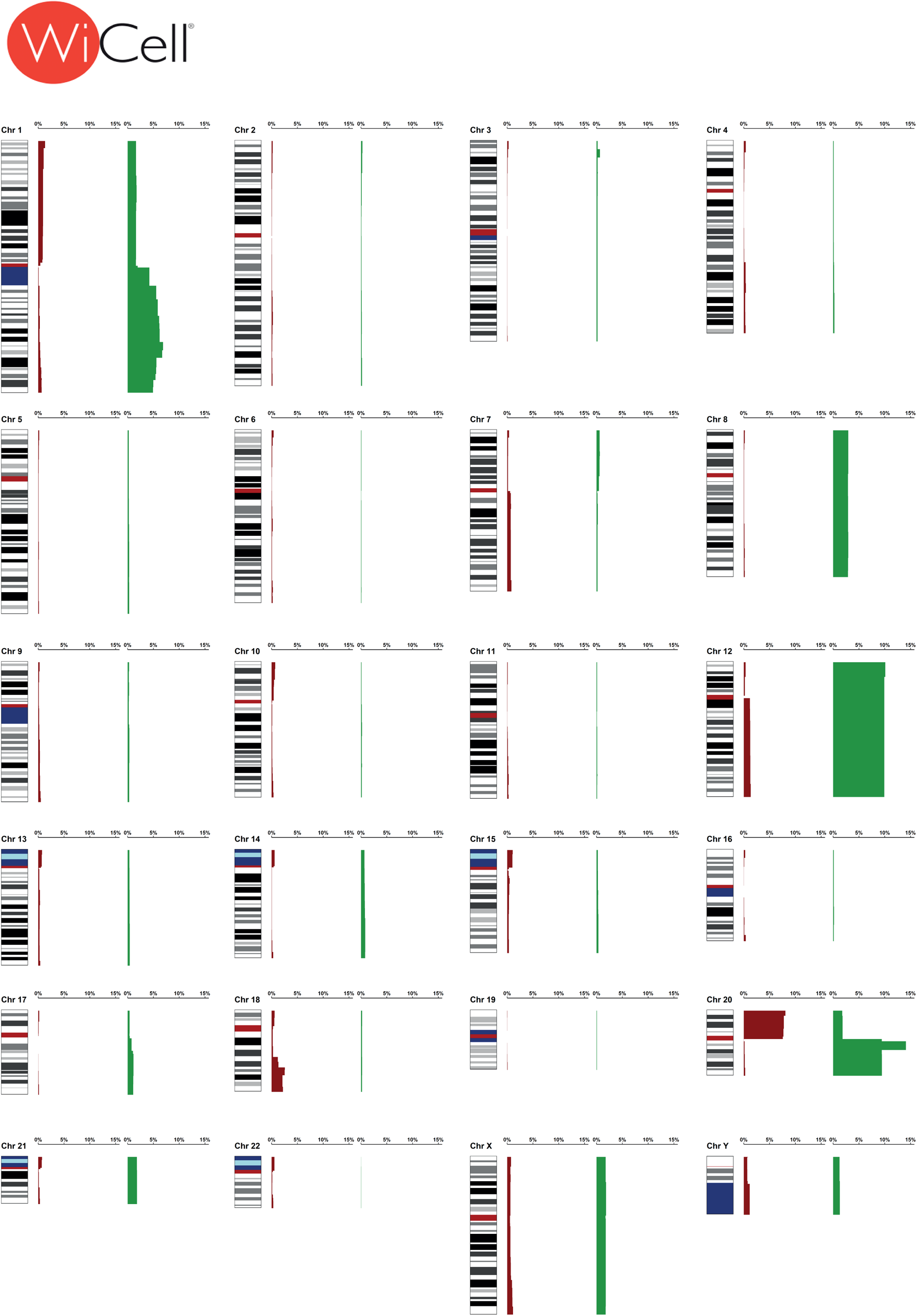
An ideogram depicting the frequency at which a particular cytoband is represented in abnormal hPSC karyotypes in the WiCell dataset. Green bars next to a chromosome represent a gain of the corresponding chromosomal region. Red bars represent losses. Related to Figure 1 and Table S2.

**Figure S2.**
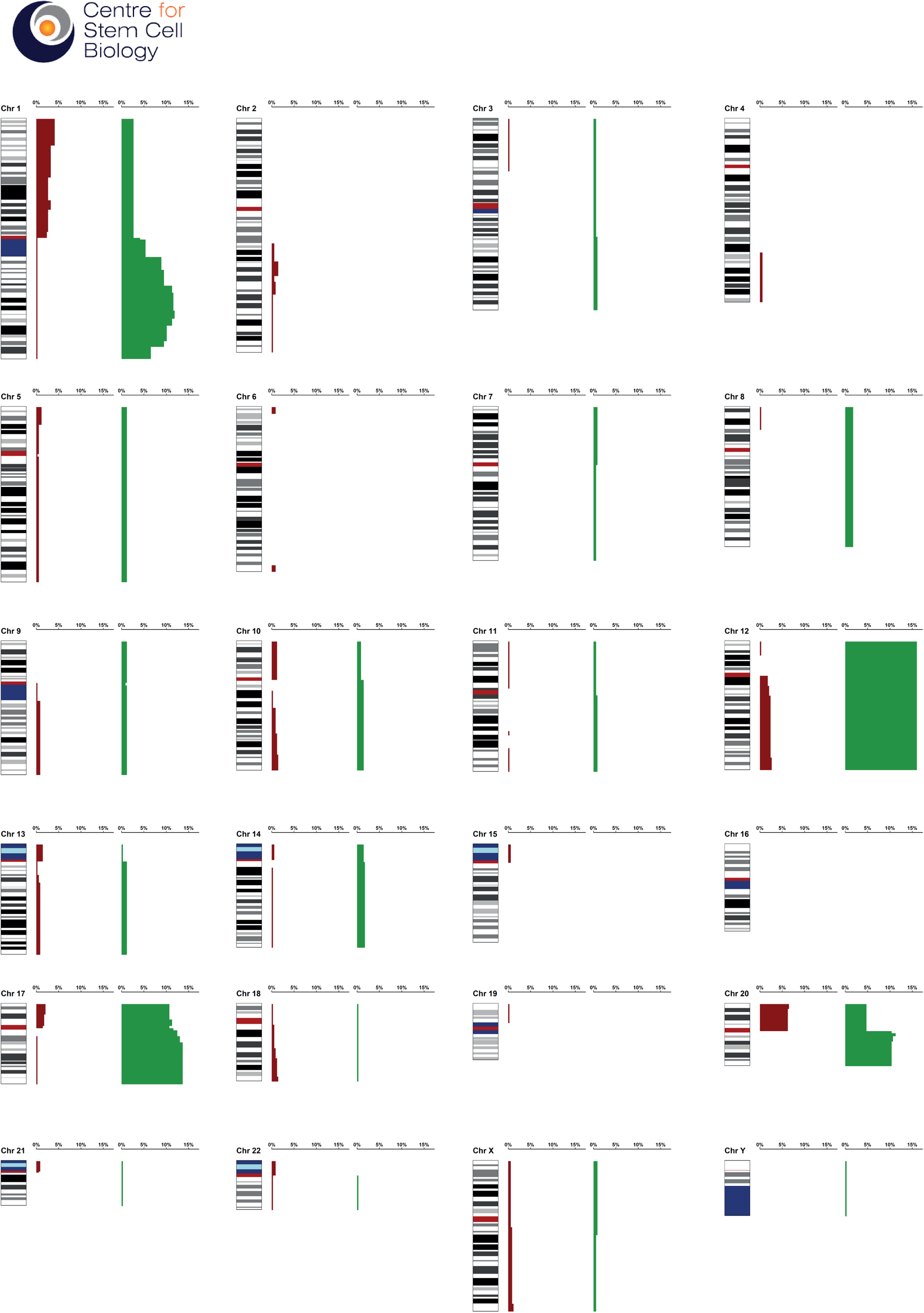
An ideogram depicting the frequency at which a particular cytoband is represented in abnormal hPSC karyotypes in the CSCB dataset. Green bars next to a chromosome represent a gain of the corresponding chromosomal region. Red bars represent losses. Related to Figure 1 and Table S2.

**Figure S3.**
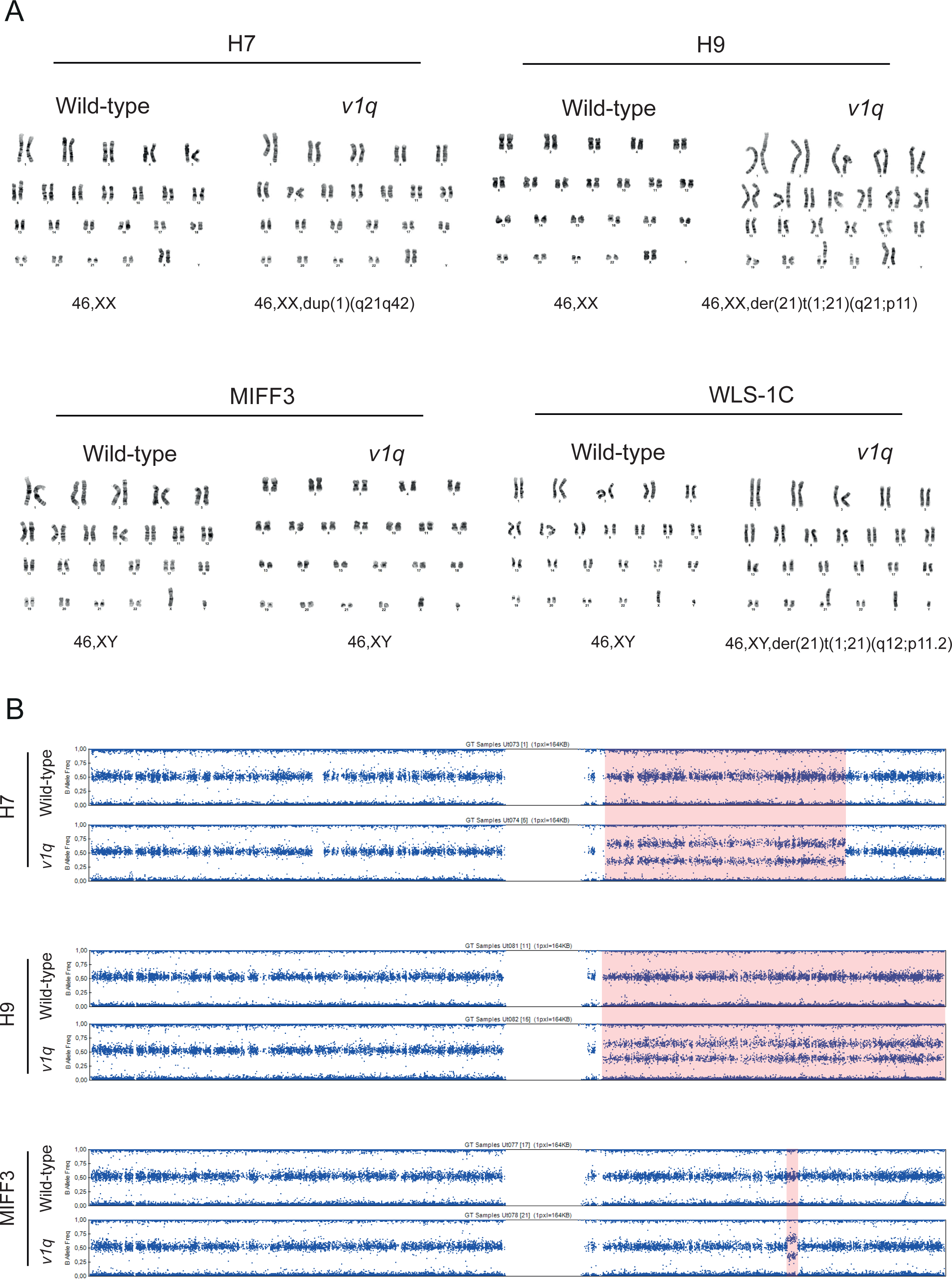
Genetic analysis of the lines used in the study. **(A)** Examples of karyotypes of H7, H9, MIFF3 and WLS-1C wild-type and *v1q* sublines. **(B)** SNP array analysis of H7, H9, MIFF3 wild-type and *v1q* sublines confirmed the presence of chromosome 1q gain. MIFF3 has a relatively small amplification of chromosome 1q that is not readily detectable by G-banding. Related to Figure 3.

**Figure S4.**
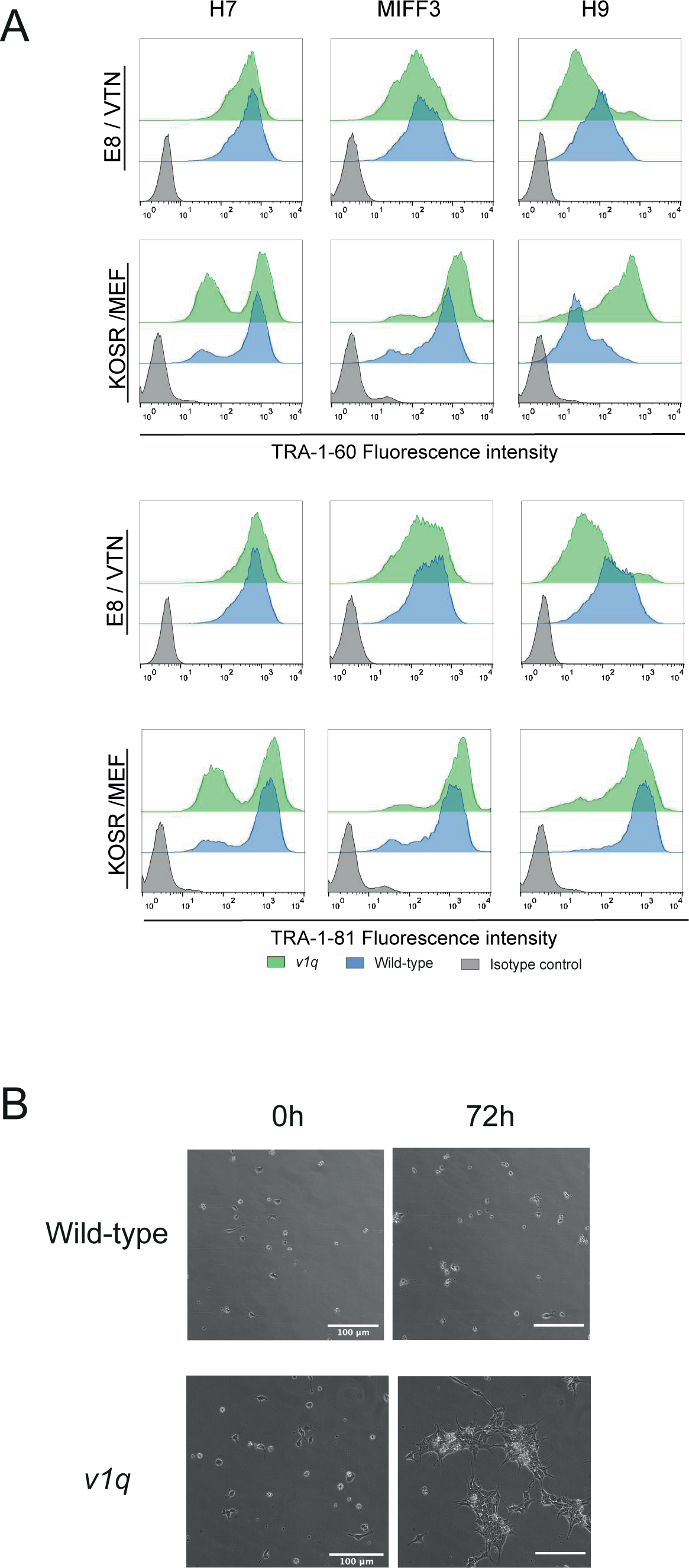
Analysis of selective advantage of *v1q* cells in E8/VTN. **(A)** Wild-type and *v1q* cells display overall similar levels of pluripotency-associated markers TRA-1-60 and TRA-1-81 in E8/VTN and in KOSR/MEF conditions. Related to Figure 3. **(B)** Frozen frames from time-lapse videos of H7 (top panels) and H7 *v1q* (bottom panel) cells. Scale bar: 100µm. Related to Figure 3.

**Figure S5.**
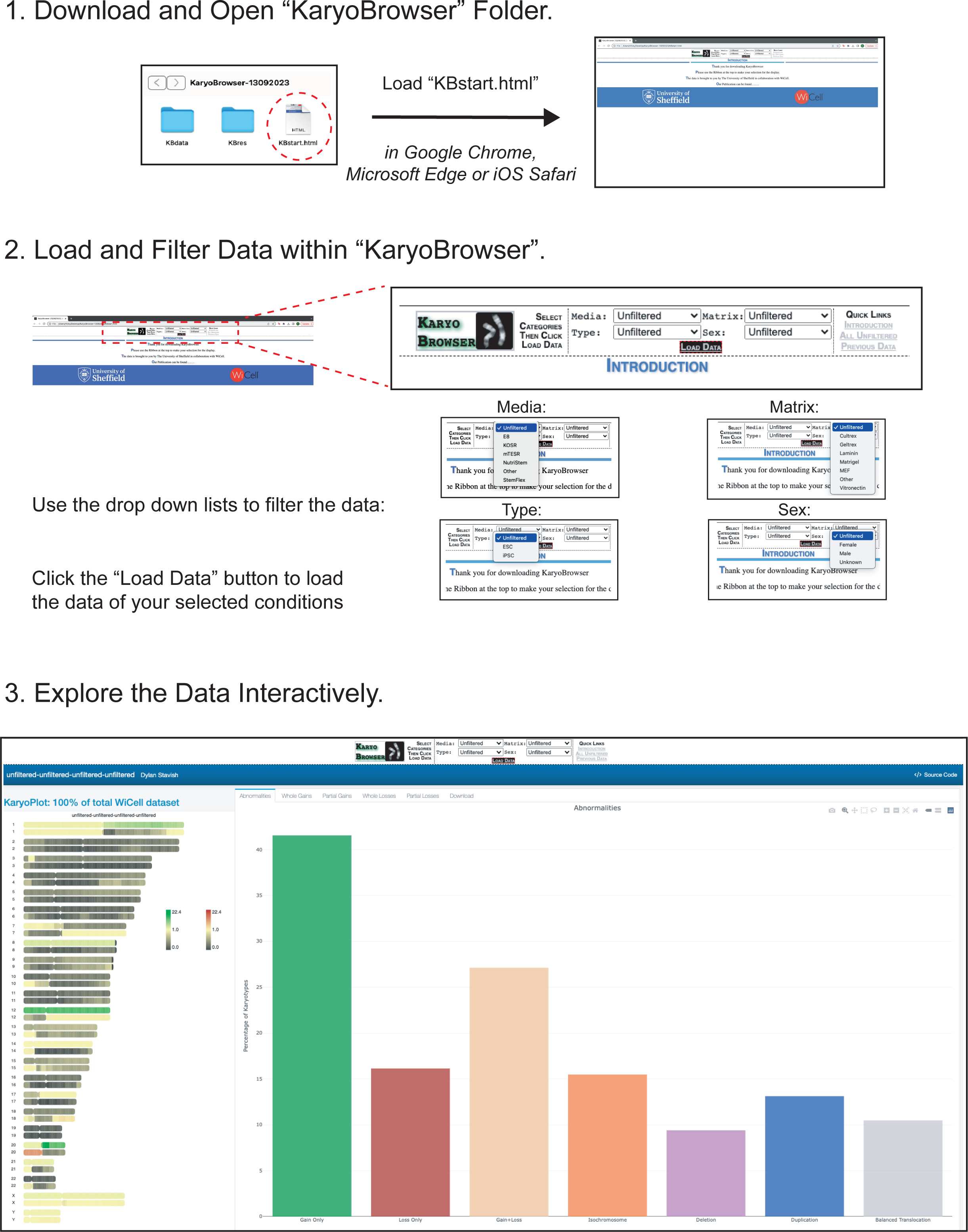
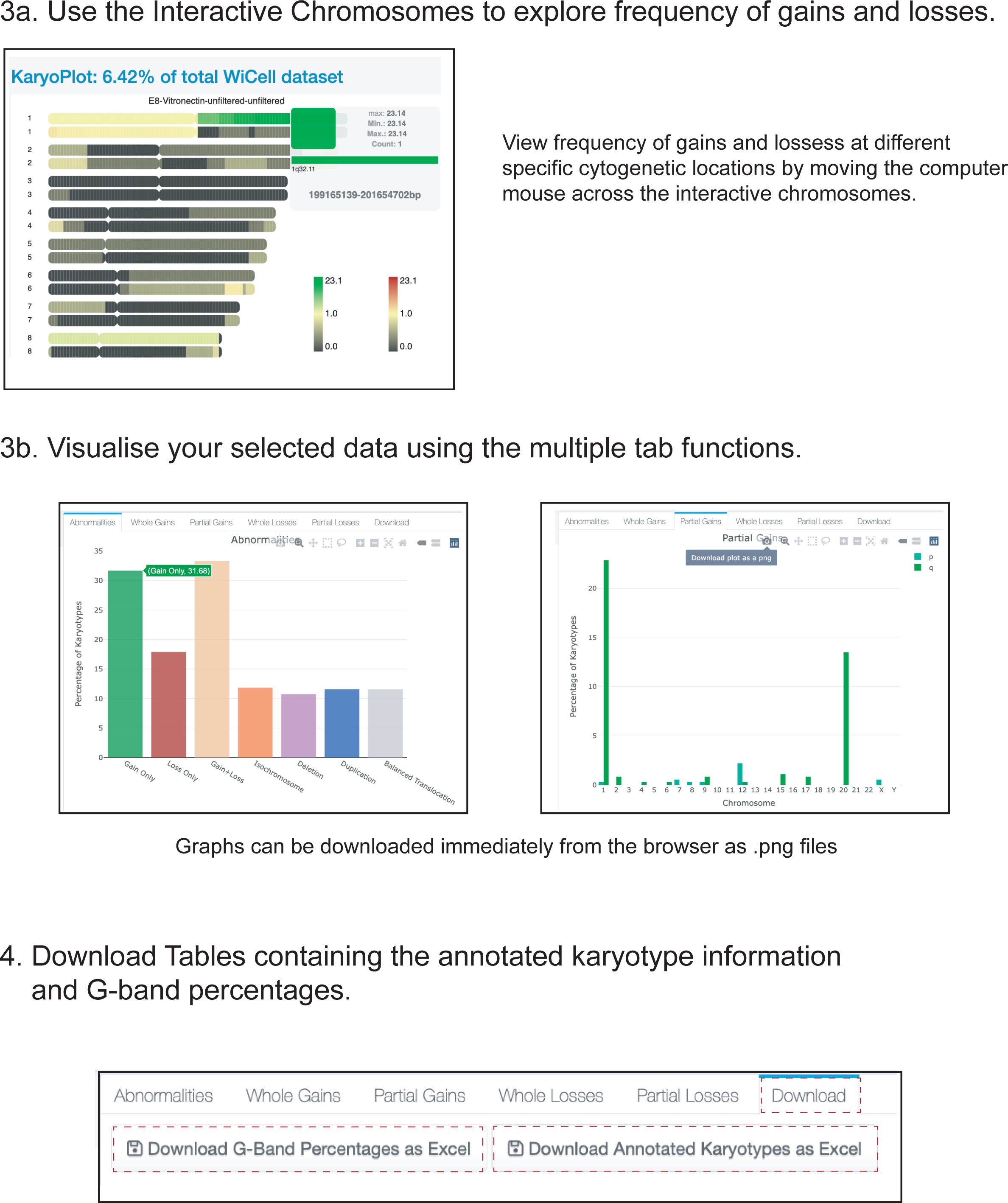
Instructions for using the interactive KaryoBrowser for exploring the karyotyping dataset collated in this study. Related to Table S1.

### Supplemental Video

**Video S1.** Time-lapse video of H7 (left) and H7 *v1q* cells (right) grown in E8/VTN. Images were taken every 10 min over 72h.

**Supplemental Tables** are available on request.

## MATERIALS AND METHODS

### Human pluripotent stem cell (hPSC) lines

Wild-type hPSCs used in this study were H7 (WA07)^14^, H9 (WA09)^14^, MIFF-3^27^ and WLS-1C^28^. Wild-type sublines were karyotypically normal (based on at least 20 metaphases analysed by G-banding of cell banks prior to experiments and at various time points upon subsequent passaging) and did not possess a commonly gained 20q11.21 copy number variant (as determined by quantitative PCR for copy number changes^11,29^ and/or Fluorescent In Situ Hybridisation^11^).

Genetically variant hPSCs used in this study and their karyotypes were: H7 *v1q* [46,XX,dup(1)(q21q42)] (20 metaphases analysed), H9 *v1q* [46,XX,der(21)t(1;21)(q21;p11)] (20 metaphases analysed), MIFF-3 *v1q* [46,XY] (20 metaphases analysed) and WLS-1C *v1q* [46,XY,der(21)t(1;21)(q12;p11.2) (20 metaphases analysed). Of note, although MIFF-3 *v1q* appeared diploid by G-banding analysis, a duplication of q32.1 on chromosome 1 was detected by qPCR and confirmed by SNParray analysis. The H7 *v1q* cell line and the fluorescently labelled H7 subline, H7-RFP, were established and described previously^30^. The WLS-1C *v1q* cells were kindly provided by STEMCELL Technologies. H9 *v1q* and MIFF-3 *v1q* were established in this study by cloning out spontaneously arising variants from mosaic cultures using single cell deposition by fluorescent activated cell sorting. Single cells from mosaic cultures were sorted directly into individual wells of a 96 well plate using a BD FACS Jazz (BD Biosciences) and cultured to form colonies over 2-3 weeks. The resulting colonies were expanded in culture and subsequently frozen to establish cell banks. At the time of freezing, sister flasks were sent for either karyotyping by G-banding and/or SNP array profiling and assessment of the relative copy number of commonly identified genetic changes by qPCR^11,29^. To facilitate cell competition assays, fluorescently labelled sublines of wild-type H9 and MIFF-3 and *v1q* lines were generated, as previously described^30^. Briefly, cells were transfected with pCAG-H2B-RFP plasmid (a kind gift from Dr Jie Na, Tsinghua University, Beijing) or pCAG-H2B-GFP plasmid^31^ (Addgene, cat. no. 184777) using Neon Transfection System (Cat. # MPK10025; Thermo Fisher Scientific). The stably transfected cells were selected by growing in medium supplemented with puromycin (Cat. # A11138; Thermo Fisher Scientific), bulk-sorted to enrich for RFP or GFP-expressing cells and subjected to karyotyping by G-banding and/or SNParray and qPCR analysis for the presence of 1q gain.

### Karyotyping database assembly and analyses

Karyotyping data analysed in this study was collected by WiCell (Madison, USA) and the Centre for Stem Cell Biology (CSCB) (Sheffield, UK) over a period of 2009-2021 and 2002-2019, respectively. While CSCB data was mainly generated from hPSC lines grown in-house, WiCell data contained hPSC samples that were grown not only in-house but also submitted to WiCell for cytogenetic analysis by different laboratories. The WiCell dataset builds on the dataset reported in^10^ (∼1,700 karyotypes) and adds a further 10 years of data collection (∼18,000 karyotypes), which equates to more than a 10-fold increase in cytogenetic data analysed.

The data was stored in a format of standard International System for Human Cytogenetic Nomenclature (ISCN), with some associated data, such as the data of submission, to the sample. WiCell data also contained media/matrix information according to customer submission. This information was screened, and blanket terms were assigned to both media and matrix for the purposes of tracking changes related to these variables but also to respect the anonymity of the WiCell customers. The CSCB dataset was curated to remove repeated sampling of the same cell line, especially given the CSCB’s active work using genetically variant hPSC lines. This same curation was not possible for the WiCell datasets due to the anonymised data. Errors in the karyotype nomenclature were corrected where possible and speculative changes denoted with question marks (?) were treated as correct calls. The karyotypes for mosaic cultures were split up for classification of karyotypic changes but kept together when assessing the number of cultures with abnormal karyotypes. In assessing the matrices for growing the cells Cultrex, which had only a ∼12% abnormality rate, was omitted from the matrix comparison as there was evident sampling bias with ∼94% of karyotypes of cells grown on Cultrex coming from just one institute.

Once curated, the data was run through CytoGPS^15^ (http://cytogps.org/) or through their offline package (https://github.com/i2-wustl/CytoGPS) which can read written karyotypes and translate them into information of losses and gains for chromosomal regions. The new dataset was put into the R package, RCytoGPS^32^, for further analysis and visualisation using ggplot2 (https://ggplot2.tidyverse.org/) and magick (https://cran.r-project.org/web/packages/magick/index.html). The browser-based database was assembled using RMarkdown (https://bookdown.org/yihui/rmarkdown/) and FlexDashboard (https://pkgs.rstudio.com/flexdashboard/). CytoGPS data was used for plotting interactive ideograms using ChromoMap^33^ and bar charts using rplotly (https://plotly-r.com). A Cascading Style Sheets (CSS) structure was added to allow for browser-based selection of parameters.

### Human pluripotent stem cell culture

For E8/VTN culture conditions, hPSC were grown on vitronectin (VTN-N) (Cat. # A14700, Life Technologies)-coated flasks (5 µg/ml in Dulbecco’s phosphate buffered saline (PBS) without Calcium and Magnesium) in modified E8 medium^34^ prepared in house, consisting of DMEM/F12 (Cat. # D6421; Sigma-Aldrich) supplemented with 14 µg/l sodium selenium (Cat. # S5261; Sigma-Aldrich), 19.4 mg/l insulin (Cat. # A11382IJ; Thermo Fisher Scientific), 1383 mg/l NaHCO_3_ (Cat. # S5761; Sigma-Aldrich), 10.7 mg/l transferrin (Cat. # T0665; Sigma-Aldrich), 10 ml/l Glutamax (Cat. # 35050038; Thermo Fisher Scientific), 40µg/l FGF2-3 (Cat. # Qk053; Qkine) and 2 µg/l TGFβ1 (Cat. # 100-21; Peprotech). For time lapse experiments, E8 was prepared using DMEM/F12 without phenol red (Cat. # D6434; Sigma-Aldrich).

For KOSR/MEF culture conditions, hPSCs were grown on a layer of mitotically inactivated mouse embryonic fibroblasts (feeders) in KOSR-based medium consisting of KnockOut DMEM (Cat. # 10829018; Thermo Fisher Scientific), 20% KnockOut Serum Replacement (Cat. # 10828028 Thermo Fisher Scientific), 1x Non-essential amino acids (Cat. # 12084947; Fisher Scientific), 1mM L-Glutamine (Cat. # 25030081; Thermo Fisher Scientific), 0.1mM 2-mercaptoethanol (Cat. # 11528926; Fisher Scientific) and 8ng/ml FGF2-G3 (Cat. # Qk053; QKine).

Cells were fed daily and maintained at 37°C under a humidified atmosphere of 5% CO_2_ in air. Routine passaging was performed every 4-5 days using ReLeSR (Cat. # 05873; STEMCELL Technologies) according to manufacturer’s instructions in E8/VTN or by treatment with Collagenase IV (Cat. # MB-121-0100; Cambridge Bioscience Ltd) and manually scraping of colonies grown in KOSR/MEF.

### Karyotyping by G-banding

Karyotyping by G-banding was performed by the Sheffield Diagnostic Genetics Service (https://www.sheffieldchildrens.nhs.uk/sdgs/), as previously described^11,35^. Briefly, hPSCs were treated with 0.1 µg/ml KaryoMAX Colcemid (Cat. # 15212012; Thermo Fisher Scientific) for up to 4h. The cells were then harvested with trypsin, re-suspended in pre-warmed 0.0375M KCl hypotonic solution. After incubating for 10 min at room temperature, cells were pelleted and resuspended in fixative (3:1 methanol:acetic acid). Metaphase spreads were prepared on glass microscope slides and G-banded by brief exposure to trypsin and stained with 4:1 Gurr’s/Leishmann’s stain (Sigma-Aldrich). Slides were scanned, metaphase images captured and analysed using the Leica Biosystems Cytovision Image Analysis system (version 7.3.2 build 35).

### SNP arrays

SNP array profiling was performed on Infinium Global Screening Array-24 v3.0 BeadChip GSA. Array data was obtained from the HuGe-F as a Genome Studio vs. 2.0.4 (Illumina, Eindhoven, The Netherlands) project using the hg38 reference genome, as described previously^36^.

### Competition assays

To assess selective advantage of variants and MDM4 overexpressing cells, mixing experiments were performed using matched fluorescently labelled and unlabelled lines, as follows: H7 *v1q* with H7-RFP cells, H9 *v1q* with H9-RFP cells, MIFF3 *v1q*-H2B-GFP with MIFF3 counterparts, and MIFF3-GFP cells with MIFF3 *v1q* or MIFF3-MDM4 cells. Cells were harvested to single cells by treating with Accutase (A6964; Sigma-Aldrich) and counted. Cells were then mixed to contain 10% *v1q* cells, or MDM4 overexpressing cells, with the respective wild-type population. Cell mixes were plated into either E8/VTN or KOSR/MEF conditions with the addition of 10µM Y-27632 for 24h. Cultures were grown and passaged as described above. At each passage, one flask was kept for further passaging while a parallel flask was harvested for assessment of the ratio of individual sublines using flow cytometry.

### Clonogenic assays

Cells were harvested by treating with Accutase (A6964; Sigma-Aldrich) at 37°C for 10 min to create a single cell suspension. Cells were washed with DMEM/F12, pelleted at 200*g* and resuspended in fresh DMEM/F12. Cells were then seeded at a density of ~500 cells per cm^2^ in 24 well plates, which had been pre-coated with either vitronectin or plated with mouse embryonic fibroblasts (MEF), and incubated with 0.5ml of pre-warmed media containing 10µM ROCK inhibitor Y-27632 (Cat. # A11001-10; Generon). After two days, a fresh 0.5ml of media containing 10µM ROCK inhibitor Y-27632 was added on top of the existing media. After four days, the media was fully replaced with 0.5ml of media without ROCK inhibitor. On day 5, cells were fixed using 4% PFA and stained for NANOG (Cat. # 4903; Cell Signalling Technology) and nuclei were stained with Hoechst 33342. Images were taken on the InCell 2200 and tiled across the well. The tiled images were stitched together for analysis and colonies were counted from images using a CellProfiler^37^ pipeline.

### Growth curve analysis

Initial assessment of population growth of wild-type and *v1q* cells was done using MIFF3 H2B-GFP and MIFF3 *v1q* H2B-GFP cells plated at 30,000 cells per cm^2^ in 96 well plates. Cells were plated directly into test conditions, i.e., E8/VTN or KOSR/MEF with 10µM Y-27632 (Cat. # A11001-10; Generon). After 24 hours, the media was replaced and Y-27632 was removed. Cells were imaged for GFP signal at day 0, 1, 2 and 3 using the InCell Analyzer with 16 set positions in each well. The GFP signal at each position was calculated as area of image covered and tracked over the days. The resulting average of the fields was used to calculate the fold change in growth.

To assess the effect of KOSR addition to E8, MIFF3 cells and MIFF3 *v1q* cells growth rate analysis was performed, as previously described^7^. In brief, cells were plated at 30,000 cells per cm^2^ in 96-well plates. Cells were plated directly into test conditions, i.e., E8/VTN, KOSR/MEF or E8+20% KOSR/VTN with 10µM Y-27632 (Cat. # A11001-10; Generon). After 24 hours, Y-27632 was removed and the media replenished daily. Plates were fixed daily with 4% PFA and nuclei counterstained with Hoechst 33342 prior to imaging on InCell Analyzer. The resulting images were analysed using a CellProfiler^11^ pipeline to calculate cell numbers.

### Time-lapse analysis

Time-lapse microscopy was performed at 37°C and 5% CO_2_ using a Nikon Biostation CT. To perform lineage analysis, cells were imaged every 10 min for 72 h using 10x air objective. Image stacks were compiled in CL Quant (Nikon) and exported to FIJI (Image J)^38^ for analysis. Lineage trees were constructed manually from FIJI movies. Individual cells were identified in the first frame and then tracked in each subsequent frame until their death, division or the end of the movie. The timing of cell death or division for each cell was noted and then used to reconstruct lineage trees of founder cells using Interactive Tree Of Life (iTOL)^39^ software.

### Flow cytometry

To assess the ratio of individual sublines in mixing experiments, cells were harvested with TrypLE (Cat. # 11528856; Thermo Fisher Scientific) or Accutase (A6964; Sigma-Aldrich) and resuspended in DMEM/F12. The sample was then pelleted by centrifugation at 200 *g* for 5 min and subsequently fixed with 4% PFA for 10 min. The sample was washed with PBS and stored at 4°C until analysis. Samples were run on the BD FACS Jazz (BD Biosciences) flow cytometer to assess ratios of unlabelled to labelled. Gates were set based on 100% labelled and 100% unlabelled samples.

For analysis of pluripotency-associated surface antigens, cells were harvested with TrypLE (Cat. # 11528856; Thermo Fisher Scientific) and resuspended in PBS supplemented with 10% Foetal Calf Serum (FCS) at 1×10^7^ cells/mL. Primary antibodies SSEA-3^40^, TRA-1-81^41^ and TRA-1-60^41^, prepared in-house as described previously^42,43^, were added to cells suspension and incubated for 30 min at 4°C. After washing with PBS supplemented with 10% FCS, cells were incubated with secondary antibody (Goat anti-Mouse AffiniPure IgG+IgM (H+L), Cat. # 115-605-044-JIR; Stratech) at 1:200 for 30 min at 4°C in the dark. After washing twice with PBS supplemented with 10% FCS, analysed on BD FACS Jazz (BD Biosciences). Baseline fluorescence was set using the isotype control antibody P3X, an antibody secreted from the parental myeloma cell line P3X6Ag8^44^.

Flow cytometry for cleaved caspase-3 was performed to assess levels of apoptotic cells in cultures, as previously described30,31. In brief, the old media, containing apoptotic cells which had detached from the flask, was collected into a 15ml Falcon tube. The remaining cells within the flask were harvested with TrypLE (Cat. # 11528856; Thermo Fisher Scientific) and added to the 15ml tube containing the collected culture medium and apoptotic cells. The collated sample was pelleted by centrifugation at 270 *g* for 5 min and subsequently fixed with 4% PFA. Cells were permeabilised with 0.5% Triton X-100 in PBS and then incubated with anti-cleaved caspase-3 primary antibody (Cat. # 9661; Cell Signalling Technology) in blocking buffer (1% BSA and 0.3% Triton X-100 in PBS). Samples were gently agitated for 1h at room temperature or overnight at 4°C, prior to washing and staining with secondary antibody (Goat anti-Rabbit AffiniPure IgG+IgM (H+L), Cat. # 111-605-003-JIR; Stratech) for 1h at room temperature in the dark. Cells were then washed and analysed on BD FACS Jazz (BD Biosciences). Baseline fluorescence was set using secondary antibody-only stained samples.

### Western blotting

Cells were lysed in 1x RIPA Buffer pre-warmed to 95°C and the total protein concentration was normalised using the Pierce BCA Protein Assay (Cat. # 23227; Thermo Fisher Scientific). Proteins (20 µg, 10µg and 5 µg /sample) were resolved by SDS-PAGE on Mini-PROTEAN TGX Stain-Free Gels (Cat. # 4568086; Bio-Rad) and were run alongside a Page Ruler prestained protein ladder (Cat. # 26616; Thermo Fisher Scientific). Proteins were then transferred onto a 0.2µm PVDF membrane (Cat. # 1704156; Bio-Rad) using a Trans-Blot Turbo Transfer System (Bio-Rad). The membrane was blocked in 5% milk for one hour, washed three times with TBS-T (50 mM Tris-HCl (pH 7.5), 150 mM NaCl, 0.1% (v/v) Tween 20) and then incubated with primary antibodies for MDM4 (Cat. # 04-1556; Sigma-Aldrich) at 1:1,000 dilution, or β-ACTIN (Cat. #60008-1-Ig; Proteintech) at 1:5,000 dilution. Following three washes with TBS-T, the membrane was incubated with secondary antibody Anti-Mouse IgG (H+L), HRP conjugate Cat. # 1706516; Bio-Rad) at 1: 5,000 dilution for 1h. After three washes, immunoreactivity was visualised using Clarity Western ECL Substrate (Cat. # 1705061, Bio-Rad) and signal captured on either x-ray film or digital detection using the LI-COR C-DiGit (LI-COR Biosciences).

### RNA extraction, sequencing and analysis

Three biological replicates of wild-type and *v1q* sublines of the H7 and H9 hPSC lines from E8/VTN cultures were used for RNA extraction and RNAseq analysis. Prior to RNA isolation, cells were lysed using Buffer RLT (Cat. # 79216; Qiagen) and stored at −80°C. RNA was isolated, libraries constructed and sequenced by GENEWIZ (Azenta Life Sciences, UK).

Briefly, libraries were prepared for Illumina (New England Biolabs, Ipswich, USA) and the library preparations were sequenced on an Illumina Novaseq platform (Illumina, San Diego, USA) to generate 150 bp paired-end reads. The sequencing reads were aligned to the GRCh38 human reference genome using STAR aligner v.2.5.2b. Unique gene hit counts were calculated using featureCounts from the Subread package v.1.5.2 and were normalized into transcript per million mapped reads (TPM), based on the length of the gene and reads count mapped to it. Differential gene expression analysis was performed using the DESeq2 R package (1.18.0)^45^. The Wald test was used to generate p-values and log2 fold changes. Genes with an adjusted p value of <0.05 and absolute log2 fold change >1 were considered differentially expressed per comparison. Enrichment analysis of differentially expressed genes was performed using the gene set enrichment analysis (GSEA) functions^46^.

### MDM4 siRNA knock-down

To knockdown the expression level of MDM4 in MIFF-3 *v1q* cells, we used MISSION esiRNA for MDM4 (ESIRNA HUMAN MDMX, Cat. # EHU005381-20UG; Sigma-Aldrich) and MISSION esiRNA for Renilla Luciferase as a control (ESIRNA RLUC Cat. # EHURLUC-20UG; Sigma-Aldrich). A 500µl transfection reaction included 50 nM siRNA and 5.6 µl DharmaFECT 1 Transfection Reagent (Cat. # T-2001-03; Horizon Discovery Ltd) in Opti-MEM I Reduced Serum Medium (Cat. # 10149832; Thermo Fisher Scientific). The reactions were incubated for 30 min at room temperature before mixing with 400,000 *v1q* cells in mTeSR (Cat. # 85850; STEMCELL Technologies) supplemented with 10μM Y-27632 (Cat. # A11001-10; Generon). Cells were plated into one well of 6-well plate per siRNA condition. After 18 hours the siRNA was removed, and cells were dissociated for plating into clonogenic assays. The remaining cells were pelleted and stored for qPCR analysis, as described below.

### Quantitative PCR (qPCR)

Expression of MDM4 in siRNA knockdown experiments was assessed using qPCR. RNA was isolated using a Qiagen RNAeasy Plus Mini Kit (Cat. # 74134; Qiagen), and the RNA concentration and purity determined using a NanoPhotometer (Implen, Munich, Germany). cDNA was synthesised using a high-capacity reverse transcription kit (Cat. # 4368814; Thermo Fisher Scientific). qPCR reactions were set up in triplicate, with each 10µl PCR reaction containing 1X PowerTrack SYBR Green Master Mix (Cat. # A46110; Thermo Fisher Scientific), 10µM Forward and Reverse Primers (Integrated DNA Technologies) and 10ng of cDNA. PCR reactions were run on a QuantStudio 12K Flex Thermocycler (Cat. # 4471087; Life Technologies). Following the first two steps of heating the samples to 50°C for 2 min and denaturing them at 95°C for 10 min, reactions were subjected to 40 cycles of 95°C for 15 s and 60°C for 1 min. The Ct values were obtained from the QuantStudio 12K Flex Software with auto baseline settings. Data was normalised to control GAPDH and 2^−ddCt^ calculated for relative expression in comparison to non-transfected *v1q* cells.

### MDM4 overexpression

The pCAG-MDM4 expression vector was established by cloning of MDM4 sequence into a pCAG vector^47^ containing a multiple cloning site (pCAG-MCS) using In-Fusion® Snap Assembly Starter Bundle kit (#638945; Takara Bio). A single restriction digest was performed on the pCAG-MCS vector using XhoI (Cat. # 0146, New England Biolabs) to linearize plasmid. The MDM4 sequence in pLVX-TetOne-Puro-MDM4, kindly gifted to us by Jason Sheltzer (Cat. # 195140; Addgene), was PCR amplified and the resulting fragment cloned into the linearized pCAG-MCS vector as per manufacturer’s instructions.

To generate the MIFF3 wild-type MDM4 overexpressing line, cells were transfected using the Neon Transfection System as described previously^30,31^. In brief, MIFF3 cells were dissociated to single cells using TrypLE and resuspended at 2,0×10^4^ cells/ml in “R buffer”. Transfection was performed with 5µg of plasmid DNA using 1 pulse of 1600V, 20msec width. After electroporation, the cells were immediately transferred to a vitronectin-coated 60mm diameter culture dish (Cat. # 150288; Thermo Fisher Scientific) containing E8 media supplemented with ^29^transfection cells were subjected to puromycin (Cat. # A11138; Thermo Fisher Scientific) drug selection. The cells were then expanded in the presence of puromycin selection and subsequently frozen to establish cell banks. At the time of freezing, cells from sister flasks were karyotyped by G-banding and assessed for the relative copy numbers of commonly identified genetic changes by qPCR^11,29^. Upon defrosting and subsequent culture, cells were also regularly genotyped by karyotyping and screened for common genetic changes by quantitative PCR^11,29^.

## QUANTIFICATION AND STATISTICAL ANALYSIS

Statistical analysis of the data presented was performed using either GraphPad Prism version 9.0.2, GraphPad Software, La Jolla California USA, www.graphpad.com or Real Statistics Resource Pack for Excel, Charles Zaiontz, www.real-statistics.com. Differences were tested by statistical tests as indicated in figure legends.

